# Discovery of synapse-specific RNA N6-methyladenosine readers associated with the consolidation of fear extinction memory

**DOI:** 10.1101/2022.06.07.495230

**Authors:** Sachithrani U. Madugalle, Wei-Siang Liau, Qiongyi Zhao, Xiang Li, Hao Gong, Paul R. Marshall, Ambika Periyakaruppiah, Esmi L. Zajackowski, Laura J. Leighton, Haobin Ren, Mason Musgrove, Joshua Davies, Simone Rauch, Chuan He, Bryan C. Dickinson, Lee Fletcher, Barbora Fulopova, Stephen R. Williams, Robert C. Spitale, Timothy W. Bredy

## Abstract

The RNA modification N^6^-methyladenosine (m^6^A) is critically involved in the regulation of gene activity underlying experience-dependent plasticity, and is necessary for the functional interplay between RNA and RNA binding proteins (RBPs) in the nucleus. However, the complete repertoire of m^6^A-modified RNA interacting RBPs in the synaptic compartment, and whether they are involved in fear extinction, have yet to be revealed. Using RNA immunoprecipitation followed by mass spectrometry, we discovered 12 novel, synapse-specific, learning-induced m^6^A readers in the medial prefrontal cortex of male C57/B6 mice. m^6^A RNA-sequencing also revealed a unique population of learning-related m^6^A-modified RNAs at the synapse, which includes a variant of the long non-coding RNA (lncRNA) metastasis associated lung adenocarcinoma transcript 1 (*Malat1*). m^6^A-modified *Malat1* binds to a subset of novel m^6^A readers, including cytoplasmic FMR1 interacting protein 2 (CYFIP2) and dihydropyrimidase-related protein 2 (DPYSL2) and a cell-type-specific, state-dependent, and synapse-specific reduction in m^6^A-modified *Malat1* disrupts the interaction between *Malat1* and DPYSL2 and impairs fear extinction. The consolidation of fear-extinction memory therefore relies on an interaction between m^6^A-modified *Malat1* and select RBPs in the synaptic compartment.

## Main

The extinction of conditioned fear, the reduction in responding to a feared cue, which occurs when the cue is repeatedly presented without any adverse consequence, is an evolutionarily conserved behavioural adaptation that is critical for survival. Like other forms of learning, long-lasting memory for fear extinction depends on coordinated changes in gene expression, particularly in the infralimbic prefrontal cortex (ILPFC) [1] [2,3]. We and others have shown that this process involves a variety of RNA-based mechanisms, including the activity of microRNAs [4] as well as the localised regulation of transcription by long noncoding RNAs [5] [6] [7]. However, there other regulatory mechanisms that can influence RNA metabolism and the capacity of neurons to adapt in response to learning. In particular, the RNA modification N6-methyladenosione (m^6^A) is dynamic and reversible [8] and regulates many aspects of state-dependent RNA metabolism in the central nervous system, including RNA stability, localisation, and translation [9] [10] [11]. This epitranscriptomic mark is enriched in the adult brain [12], directly involved in the acquisition of fear-related learning and memory [13], spatial memory consolidation [14] [15] and involved in dopaminergic signalling [16] [17] [18]. In addition, a population of synapse-enriched m^6^A-modified RNA has recently been discovered in the adult prefrontal cortex [19]; however, the functional relevance of localised m^6^A-modified RNA and its interacting partners in the context of fear extinction has not been elucidated.

Many RNA binding proteins (RBPs) recognise and interact with m^6^A-modified RNA, and several, including the YTHDF family of proteins, have been shown to be m^6^A readers [20] [21] [22] [23] [24] [25] [26]. Whether there are m^6^A readers in the synaptic compartment that support specific forms of learning and memory, including fear extinction is completely unknown. Given the increasing recognition that m^6^A-modiifed RNAs play important roles in brain function [11], we therefore considered their impact on this important learning process. Using RNA immunoprecipitation followed by mass spectrometry, we discovered 12 novel extinction learning-induced m^6^A readers that are primarily associated with in synaptic remodeling and synaptic vesicle endocytosis. This was accompanied by the detection of a unique population of learning-related m^6^A-modified RNAs at the synapse, including a variant of the long noncoding RNA (lncRNA) metastasis associated lung adenocarcinoma transcript 1 (*Malat1*). In addition, we found that the synapse-enriched m^6^A-modified *Malat1* interacts with a subset of novel m^6^A readers, including cytoplasmic FMR1 interacting protein 2 (CYFIP2) and dihydropyrimidase-related protein 2 (DPYSL2, also known as CRMP2), and that a cell-type-specific, state-dependent, and synapse-specific reduction in m^6^A-modified *Malat1* disrupts the interaction between *Malat1* and DPYSL2 and impairs fear extinction. The formation of fear extinction memory therefore relies on direct interaction between m^6^A-modified Malat1 and a select number of novel m^6^A readers in the synaptic compartment.

## Results

### Novel synaptic m^6^A readers associated with fear-extinction learning

Previous studies have identified m^6^A-modified RNA binding proteins, including YTHDF2/3, YTHDC1, ELAVL1, FMRP and IGF2BP1 [27] [24] [28] with some being expressed at the synapse [19]. However, no study has determined whether there are learning-specific m^6^A-modified RNA interacting RBPs in the synaptic compartment. Using synthetic biotinylated control and m^6^A-modified oligonucleotides, we set out to determine whether there are synaptic m^6^A readers that might be specific to fear-extinction learning. C57/BL6 mice were trained using a standard cued fear-conditioning task followed by either novel context exposure (retention control, RC) or extinction (EXT) training. Immediately after training, the mPFC was extracted and four were pooled for the isolation of synaptosomes followed by m^6^A-modified RNA-immunoprecipitation followed by mass-spectrometry (m^6^A-RIP-MS). In the synaptic fraction derived from RC-trained animals, we identified 76 proteins that interacted with the m^6^A-modified RNA, with many proteins associated with mitochondrial function (Fig. 1A). In comparison, within the synapse-enriched fraction of EXT trained animals, 61 proteins were found to interact with m^6^A-modified RNA (Fig. 1B). These proteins are functionally distinct from those in the RC group, and included proteins involved in synaptic vesicle clustering (SYN1, SYN2) and budding (SNAP91, DNM1, DNM3), neuron projection morphogenesis (DPYSL2, TUBB3, CRMP1) and translation elongation activity (EEF1A1, EEF1A2). Overall, we identified 12 synapse-specific m^6^A readers that were unique to EXT (Table 1). These included proteins associated with the synapse (CYFIP2, ADD1), and involved in synaptic vesicle endocytosis (PACSIN1, SH3GL1) and calmodulin binding (CAMKV, PLBC1).

**Figure 1:**
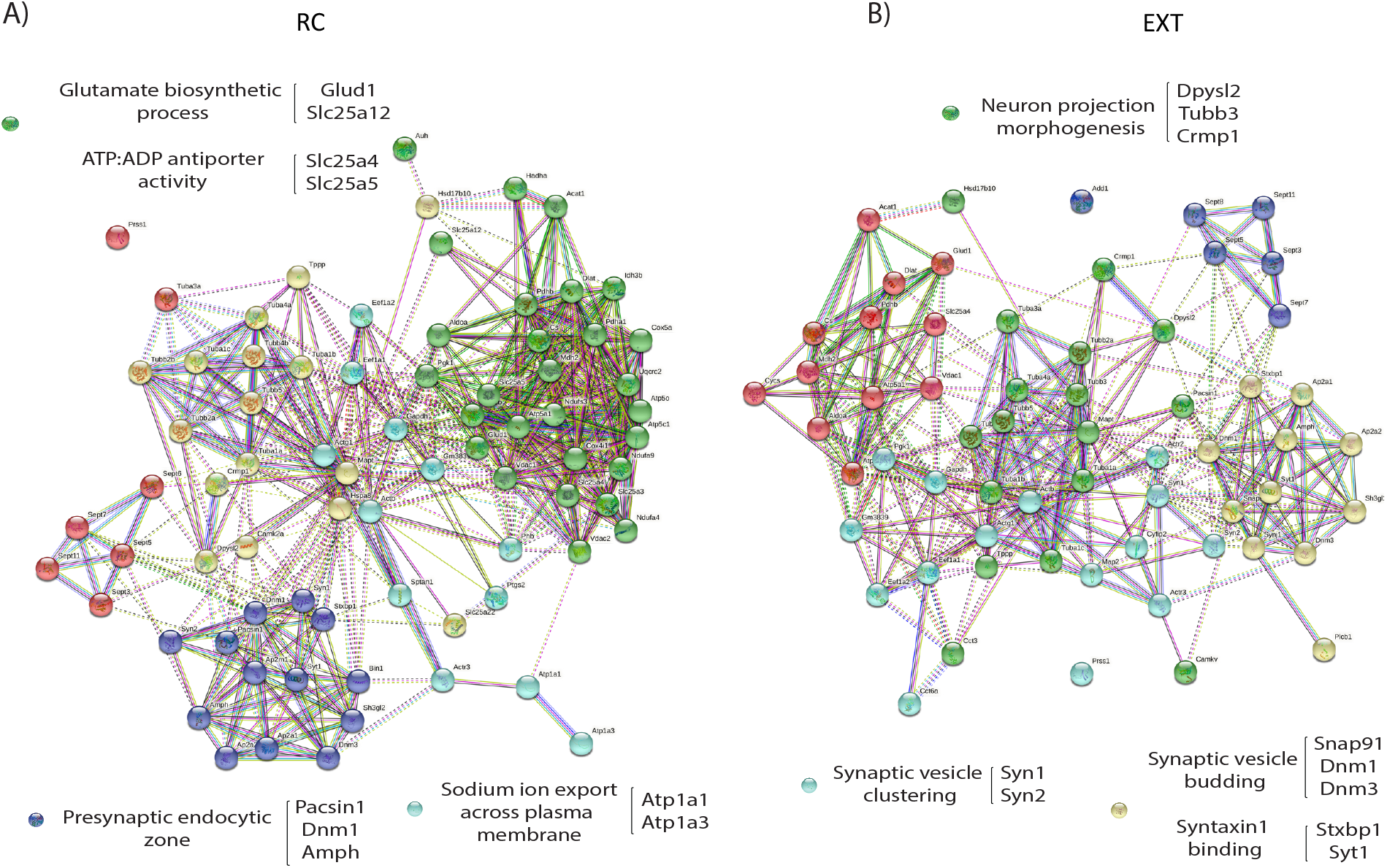
RIP-MS identifies novel synaptic m^6^A readers in the context of fear-extinction learning. A) Representative functional interaction network analysis of m^6^A interacting proteins in B) RC and C) EXT at the synapse after extinction training. RIP-MS performed using control and m^6^A synthetic biotinylated oligos to pull-down proteins in PFC synaptic lysates from retention control (RC) or extinction (EXT) trained animals (n=3 per group; each replicate is a pool of 4 PFCs).

**Table 1:** Unique m^6^A readers at the synapse after EXT learning

| Name | Function | Reference |
| --- | --- | --- |
| ADD1 | Calmodulin binding;<br>spectrin-actin network<br>assembly | [29] |
| ACTR2 | Actin polymerisation | [30] |
| CAMKV | Dendritic spine<br>maintenance, vesicle<br>formation | [31] [32] |
| CCT3 | Neurogenesis | [33] |
| CCT6A | Protein folding | [34] |
| CYCS | Apoptosis | [35] |
| CYFIP2 | Synapse function | [36] |
| PLCB1 | Neuronal proliferation | [37] |
| SEPT8 | SNARE complex formation<br>in synaptic vesicles | [38] |
| SNAP91 | Endocytosis | [39] |
| TUBB3 | Axon guidance/maintenance | [40] |
| TUBB4A | Neural process growth | [41] |

### *A* distinct population of m^6^A-modified RNA accumulates within the synaptic compartment following fear-extinction learning

Given that the pattern of RNA expression can differ based on its cellular localisation and context, we next examined the levels of m^6^A-modified RNA following fear extinction learning. tissue derived from either RC or EXT trained animals was used to isolate RNA from the synaptic compartment and subjected to low-input RNA m^6^A-sequencing. 24 libraries (input and RNA m^6^A enriched after either RC and EXT) with 3 biological replicates each (where each replicate was a pool of four PFCs) were sequenced. The peak distribution along modified transcripts in the non-synaptic fraction showed an accumulation of m^6^A in the 3’UTR (Fig. 2A, B and F). In contrast, the majority of peaks along transcripts in the synaptic fraction were found in the CDS (Fig. 2A and C). The difference in distribution of m^6^A marks in different compartments suggests the possibility that m^6^A-modified RNA serves a functionally distinct role in the synapse compared to other neuronal compartments. As such, a GO term analysis for significantly modified transcripts revealed that the non-synaptic m^6^A-modified RNA was associated with cellular transmission and nerve impulse transmission whereas the synaptic m^6^A-modified RNA was related to translation (Fig. 2D).

**Figure 2:**
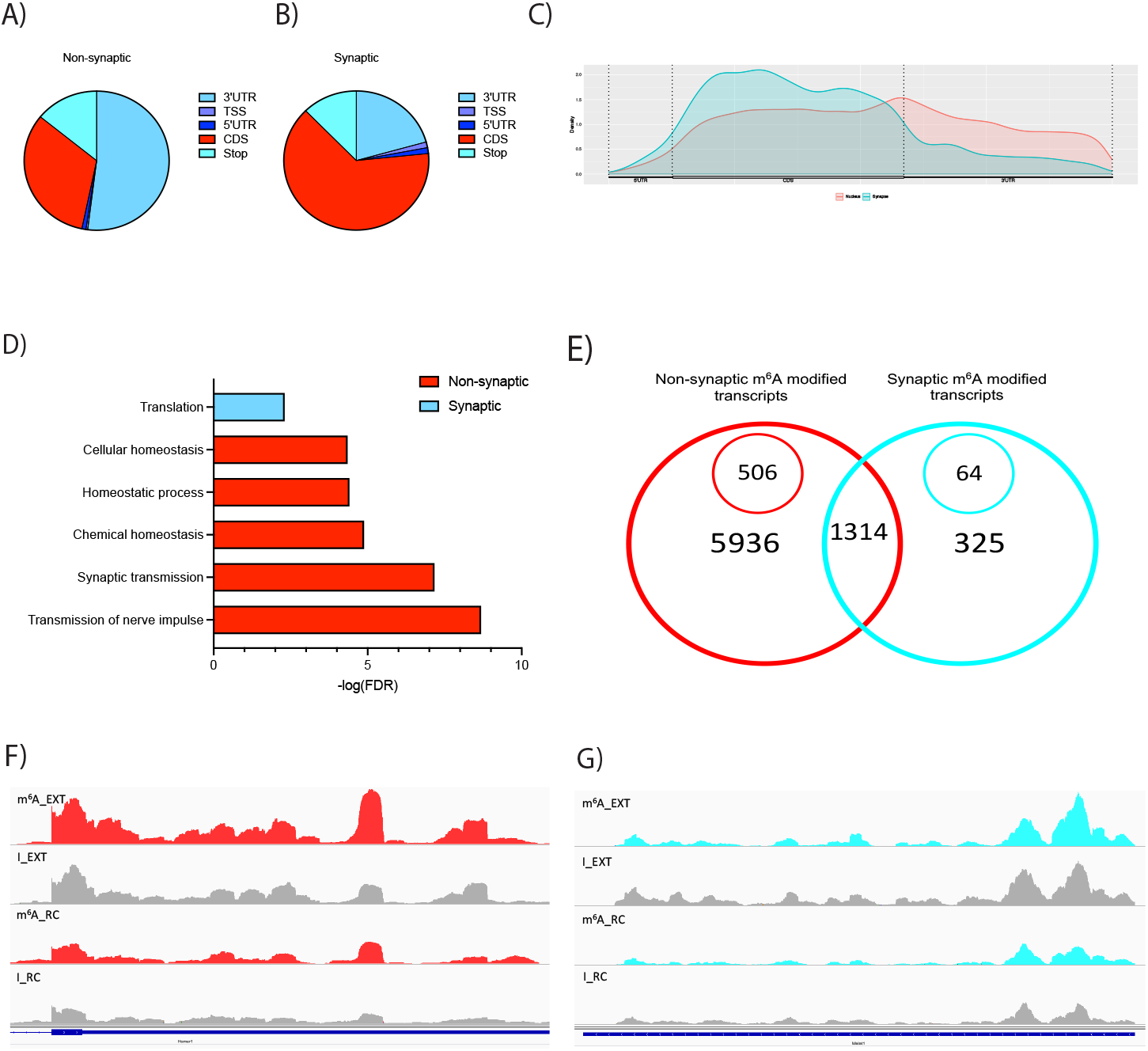
Low input m^6^A-sequencing reveals a distinct population of modified RNA at synapse after fear-extinction learning. Distribution of m^6^A marks along transcripts in (A) non-synaptic and (B) synaptic fractions show, majority of m^6^A marks in non-synaptic transcripts accumulate in the 3’UTR compared to the CDS in synaptic transcripts (C). Biological process gene ontology (GO) analysis terms for significantly modified transcripts in each fraction are shown in (D) where m^6^A-modified transcripts have different functions based on its cellular localisation. E) Venn diagram highlights 5936 m^6^A-modified transcripts in the non-synaptic fraction of which 506 are significantly modified after EXT training in comparison to input controls. Similarly, 325 synaptically localized m^6^A-modified transcripts were identified out of which 64 are significantly modified after EXT training in comparison to input controls. PFC non-synaptic and synaptic lysates from RC or EXT animals, each had input and m^6^A-enriched samples (n=3, where each replicate was a pool of 4 PFCs). Representative IGV tracks F) and G) showing *Homer1* (non-synaptic target, red) and *Malat1* (synaptic target, blue), respectively, in m^6^A and input groups after RC or EXT.

In the non-synaptic fraction, a total of 15 716 m^6^A peaks were identified, mapping to 7250 genes, where 506 transcripts were significantly enriched in comparison to input after EXT training (Fig. 2E). Many of the m^6^A modified transcripts including *Camk2a and Grin2b* [9,19] as well as *Shank1 and Homer1* [42] are known to be functionally related to plasticity and memory. In the synaptic fraction, a total of 2126 m^6^A peaks were identified, which mapped to 1639 genes. Out of these, 64 were significantly enriched in comparison to input after EXT training (Fig. 2A). In addition to several known m^6^A-modified RNAs, including *Kif5a, Atp1a2*, and ribosomal subunits *Rpl13, Rpl21, Rpl23a* and *Rpl26*, which have been previously identified at synapse under baseline conditions [19], we discovered novel, extinction learning-related m^6^A-modified transcripts, including *Trim2, Calr, Man1a2, Malat1* and *Ncam2*. Previously thought to be selectively expressed in nuclear speckles and associated with the alternative splicing of *Nlgn1* and *SynCam1* [43], *Malat1* has recently been shown to traffic to the synapse [44], suggesting that local activity of *Malat1 may* contribute to neural growth and synaptogenesis [45] [46]. Our finding that m^6^A-modified *Malat1 accumulates* at the synapse (Fig. 2G) in response to extinction learning agrees with this observation and supports the growing appreciation of the multidimensional functional capacity for lncRNAs.

### Malat1 interacts with novel m^6^A readers CYFIP2 and DPYSL2 in the synaptic compartment following fear-extinction learning

Since m^6^A-modified *Malat1* localises to the synapse in response to fear-extinction learning, we next investigated whether there are m^6^A readers that may be specific to m^6^A-modified *Malat1* at the synapse. In the RC group, 92 proteins uniquely bound to the *Malat1* oligos compared to 100 proteins in the EXT group. The proteins which interact with *Malat1* in the RC group are associated with RNA binding (EIF4A2, RPS15, RPS16), nucleotide binding (CNM1, SYN3) and glycolytic processes (ALDOA, PDK) (Fig. 3A). In comparison, EXT dependent Malat1 RBPs were associated with nucleotide binding (DDX1, ACTG1) as well as neurotransmitter secretion (STXBP1, SYN1, SYN2) and translation (RPS16, RPS20, EIF1A2). These data suggest that the proteins which bind to *Malat1* at the synapse also differ based on the learning-context.

**Figure 3:**
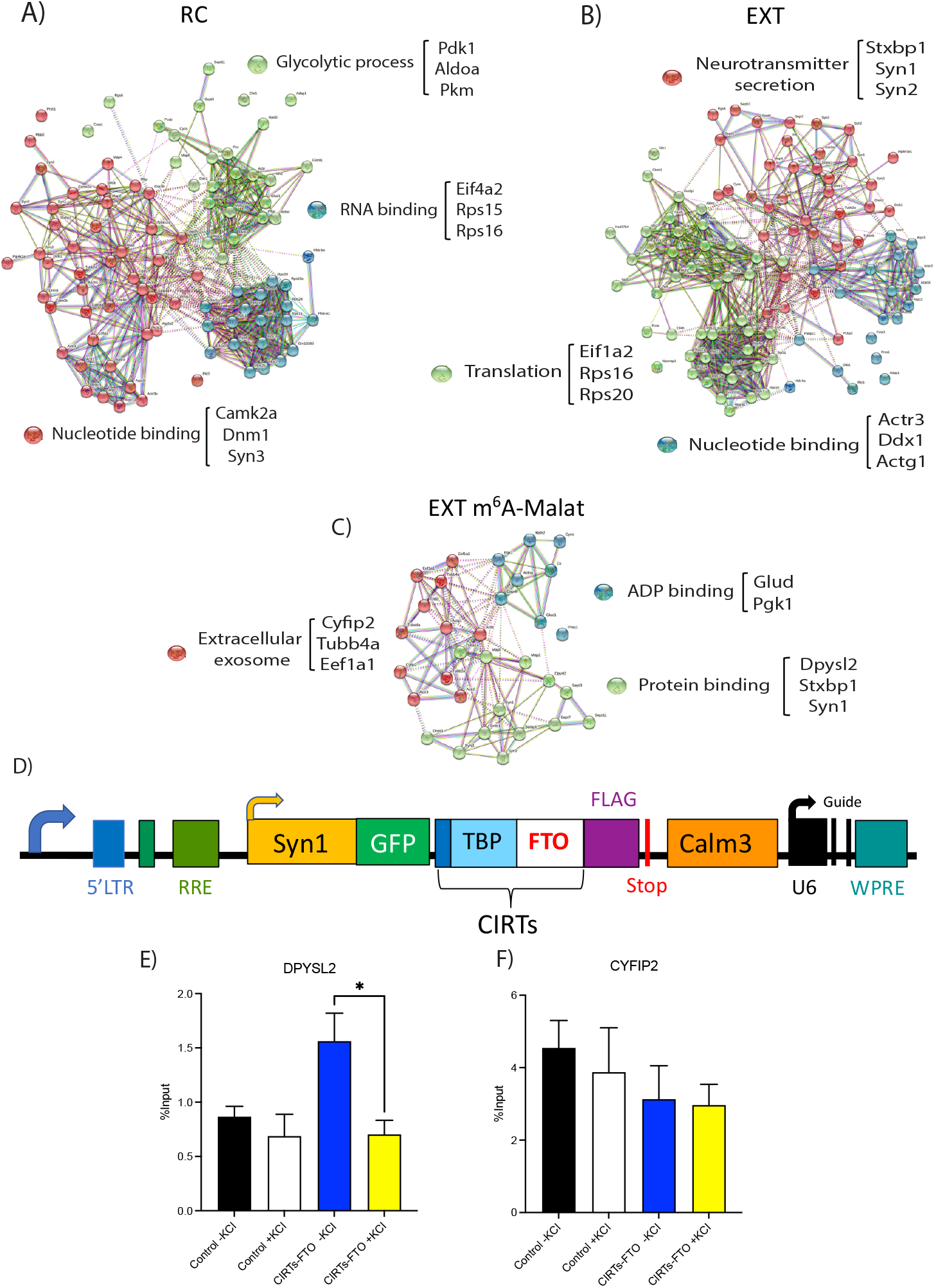
m^6^A readers involved in processes critical for memory consolidation also binds Malat1 at the synapse in the context of fear-extinction learning. Representative functional interaction network analysis of Malat1 interacting proteins in A) RC B) EXT and C) m^6^A readers which also recognize Malat1 at the synapse after extinction training (n=3 per group; each replicate is a pool of 4 PFCs). D) Schematic of CIRTs-FTO construct used for site-directed demethylation along *Malat1*. Expression levels of *Malat1* bound to E) DPYSL2 and F) CYFIP2 in control and CIRTs-FTO virus treated neurons with or without KCI treatment relative to input controls.

To determine which of the 100 EXT Malat1 proteins are also m^6^A readers, we compared them with the 62 m^6^A proteins identified at the synapse after EXT training. 32 overlapping proteins were found to recognize both m^6^A and *Malat1* at the synapse, and were associated with ADP binding (GLUD, PGK), extracellular exosomes (CYFIP2, TUBB4A, EEF1A1) and protein binding (DPYSL2, STXBP1, SYN1) (Fig. 3C). Of particular interest were the m^6^A readers CYFIP2 and DPYSL2 (also known as CRMP2) that show a strong interaction with *Malat1* at the synapse. CYFIP2 and DPYSL2 belong to the WAVE complex and serve to influence actin dynamics at the synapse and axonal growth [47] [48] [49] [50] both of which are critical for memory consolidation [51].

To confirm that the interaction is m^6^A-dependent, using an adaptation of the CIRTS system, *Malat1* was site-specifically demethylated in primary neurons by directing the m^6^A demethylase FTO to three m^6^A sites along the *Malat1* transcript (CIRTS-FTO Malat1). Upon KCI stimulation and collection of neurons, fRIP-qPCR was performed using CYFIP2 and DPYSL2 antibodies. Though the level of *Malat1* bound to CYFIP2 did not change (Figure 3D), there was a decrease in binding with DPYSL2 following *Malat1* demethylation (Figure 3E). Together, these data suggest that the interaction between m^6^A-modified *Malat1* and DPYSL2 at the synapse may be important for regulating localized activity-dependent synaptic processes underlying memory formation.

### Targeted degradation of *m^6^A-Malat1* at the synapse leads to impaired fear-extinction memory

To explore whether m^6^A-modified *Malat1* and its protein interactions at the synapse are required for fear-extinction memory, we again used the CIRTS system to trigger the degradation m^6^A-modified *Malat1* at the synapse by targeting the m^6^A reader YTHDF2 to m^6^A modified *Malat1* at the synapse. The viruses were initially validated in primary cortical neurons (Fig. 4A) followed by in vivo tests to determine whether the selective degradation of m^6^A-*Malat1* at the synapse influences fear-extinction memory.

**Figure 4:**
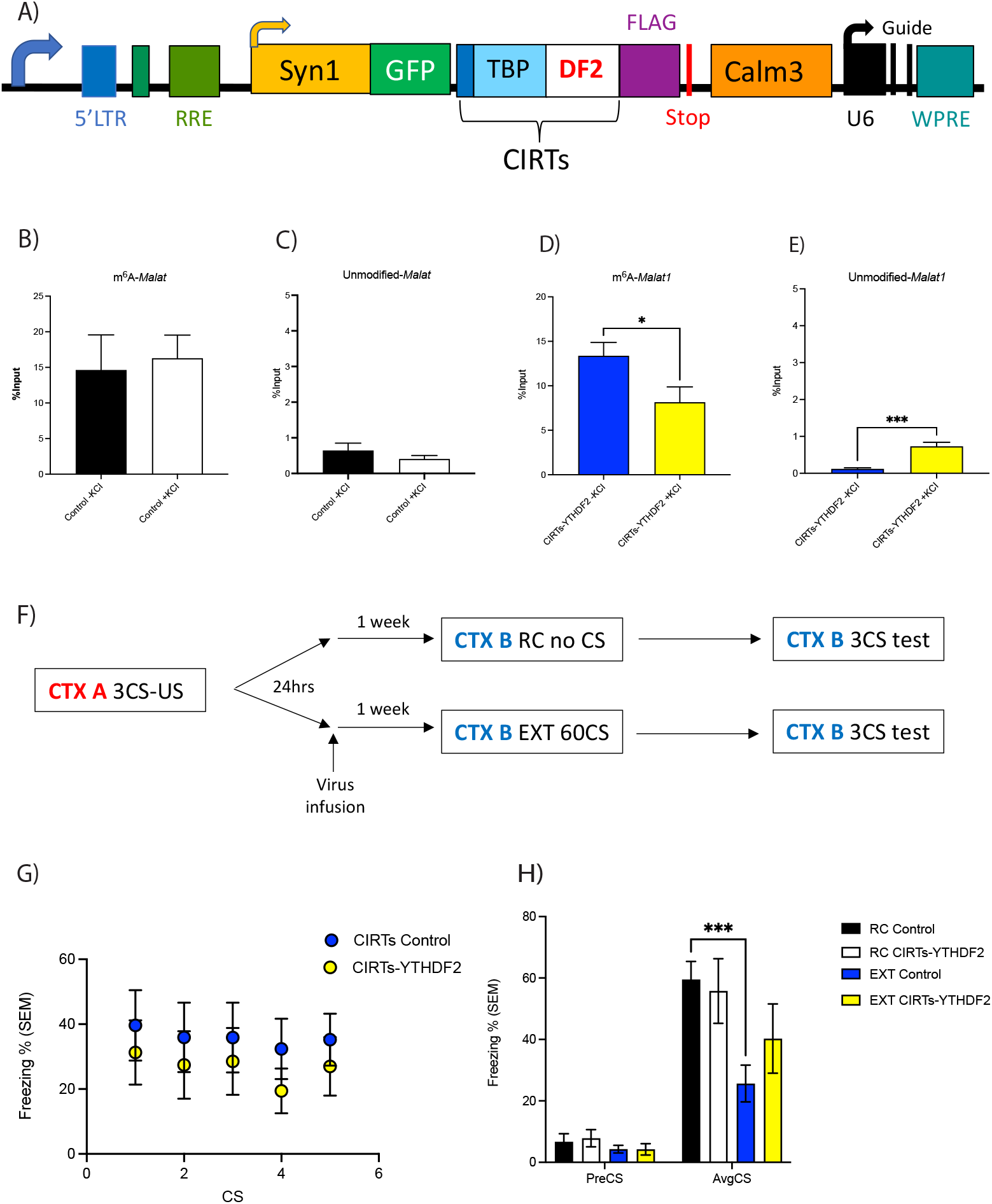
Targeted degradation of m^6^A-modified Malat1 at the synapse impairs fear-extinction memory. A) Schematic of CIRTS-YTHDF2 construct used for targeted degradation of *Malat1*. B)-E) *In vitro* validation of CIRTS-YTHDF2 construct degrading m^6^A-modified *Malat1* which is accompanied by an increase in the unmodified-*Malat1* fraction. F) Schematic of behavioural protocol used to test lentiviral mediated degradation of m^6^A-modified *Malat1* in the IL-PFC on fear-extinction memory. CTX refers to context, CS refers to conditioned stimulus (i.e. tone) and US refers to unconditioned stimulus (i.e. foot shock). G) There was no effect of CIRTS-YTHDF2 virus infusion in the IL-PFC on performance during within session extinction training. H) However, animals treated with control virus, fear conditioned and exposed to novel context (RC Control, n=9) had significantly higher freezing scores than control animals that were extinction trained (EXT Control, n=10, two-way RM ANOVA with Dunnett’s post-doc test, ***p<0.001) but not RC CIRTS-YTHDF2 animals (n=7). This effect was blocked in animals treated with CIRTS-YTHDF2 virus and extinction trained (EXT CIRTs Malat1, n=8).

Knockdown of m^6^A-modified *Malat1* prior to EXT training had no effect on within session performance (Fig. 4G). However, when testing for memory, those mice with the control virus who underwent EXT training show significantly decreased freezing scores (Fig. 4H). This same significant decrease in freezing scores is not observed in the EXT trained mice infused with the *Malat1* viruses, indicating an impairment in fear-extinction memory consolidation (Fig. 4H). Therefore, these data demonstrate that m^6^A-modified *Malat1* is critical for fear-extinction memory, and imply a role for the interaction between m^6^A-modified *Malat1* and specific RBPs in this process.

## Discussion

In this study, we have discovered that there are extinction-learning-specific m^6^A readers at the synapse. In addition, we found a unique population of synapse-enriched m^6^A-modified RNA that accumulate in response to fear-extinction learning. Fear extinction drives the expression of m^6^A-modified lncRNA *Malat1* to the synapse and promotes its interaction with novel m^6^A readers CYFIP and DPYSL2. Both proteins are critically involved in memory formation via actin polymerization at synapses and axonal growth. Consequently, upon targeted removal of the m^6^A mark on *Malat1* and degradation of m^6^A-modified *Malat1* at the synapse, RNA-protein interactions can no longer occur and fear extinction memory is impaired, suggesting a direct role for m^6^A-modified *Malat1* in the consolidation of fear extinction memory.

A significant number of novel m^6^A readers in the synaptic compartment after fear-extinction learning was revealed. These RBPs, which are a combination of indirect and direct m^6^A RBPs, include the endocytosis related protein SNAP91 [39], CAMKV and SEPTIN8 which are important for vesicle formation [32] [38], and CYFIP2, which is necessary for synapse formation [36]. Interestingly, all these processes are associated with memory consolidation, suggesting m^6^A facilitates the formation of memory through direct effects on proteins involved in various aspects of synaptic plasticity.

Many of the m^6^A-modified RNAs detected at the synapse in previous studies, such as *Homer1, Shank1* and *Camk2a* were not detected at the synapse after EXT learning, but rather in the non-synaptic fraction. This is likely due to the time point at which PFC tissue were collected. Tissue was collected immediately after EXT training; however, it is possible time after training is required for the localisation of transcripts to the synapse. In agreement with previous data, the majority detected in the non-synaptic fraction contained m^6^A peaks in the 3’UTR [52] whereas m^6^A peaks were more likely to occur in the CDS of RNAs detected in the synaptic fraction. The variation in the distribution of m^6^A peaks among transcripts suggests the functional role of m^6^A-modified RNA depends on RNA localisation. This is supported by the finding that non-synaptic transcripts were associated with cellular transmission. In contrast, m^6^A-modified synaptic transcripts were associated with translation, which confirms previous observations regarding the presences of m^6^A in the CDS [53]. The functional relevance of m^6^A is likely dictated through interactions of m^6^A-modified RNA with unique reader proteins in each subcellular compartment.

One of the most interesting synapse-enriched m^6^A-modified targets was *Malat1*. Initially, *Malat1* was discovered in cancer cells but also found to be abundant in neurons. *Malat1*, a lncRNA, has previously been demonstrated to be m^6^A-modified [54] [27] [55] [12]. This m^6^A mark is believed to change the secondary structure of *Malat1* such that proteins such as HNRNPG and HNRNPC, which are involved in splicing can bind to the lncRNA [56] [57] [58] [59]. Interestingly, *Malat1* is enriched in nuclear speckles where it recruits splicing factors to genes such as *Ngln1* and *SynCam1*. These genes are involved in synapse formation and maintenance thereby enabling an indirect influence of *Malat1* on the synaptic density of hippocampal neurons [43]. *Malat1* has also been detected in the synapse, in agreement with our findings [44] although its functional relevance was not determined.

Beyond HNRNPG and HNRNPC as m^6^A readers, we found m^6^A-modified *Malat1* interacts with numerous m^6^A reader proteins, at the synapse, in the context of fear-extinction learning. m^6^a-modified *Malat1* binds to CYFIP2 and DPYSL2. CYFIP2 protein expression at the synapse [60] contributes to actin polymerisation and therefore synaptic plasticity through changes in spine morphology [61]. In fact, reduced *Cyfip2* levels impair spine maturation in CA1 pyramidal neurons and prevents the retention of spatial memory in water maze tasks [62]. DPYSL2 has also been detected in dendritic spines [63] and is associated with axonal growth, neurotransmitter release and synaptic physiology [64]. *Crmp2* conditional knock-out mice exhibit impaired hippocampal dependent learning and memory impairments in the Y-maze test, fear-conditioning and Morris water maze test [65].

Both CYFIP2 and DPYSL2 are associated with the WAVE regulatory complex (WRC). This complex is composed of five components – CYFIP1 and/or CYFIP2, NAP1 or HEM2 or KETTE, ABI2, HSPC300 or BRICK1 and WAVE1 or SCAR [66]. Loss of function of this WRC *in vivo* or in cultured neurons results in a decrease in mature dendritic spines [50]. Also, electrophysiological recordings from hippocampal slices of WAVE1 null mice showed altered synaptic plasticity and mutant WAVE1 mice have impaired hippocampal-dependent spatial memory as evaluated by the Morris water maze test [51].

In this complex, CYFIP1 is a key regulator of neuronal actin dynamics, by regulating mRNA translation and actin polymerisation [61] [67]. In addition, CYFIP proteins also recognise FMRP, which is a known m^6^A reader, involved in the export of modified RNA [24]. However, DPYSL2 is involved in the kinesin-1 dependent transport of the WAVE1 complex. Upon DPYSL2 knockdown (KD) axonal growth is inhibited and for this DPYSL2-induced axonal growth to occur, WAVE1 is required [68]. Furthermore, KD of DPYSL2 and kinesin1 resulted in the delocalisation of the WAVE1 complex from growth cones of axons. Thus, suggesting DPYSL2 mediated localisation of the WAVE1 is critical for axon formation [49]. As it is known that lncRNAs bind to proteins to promote their localisation [69], it is likely the m^6^A mark on *Malat1* acts as beacon for the binding of WAVE complex proteins. Once bound, *Malat1* may localise these proteins to the same location, to form the complex and ultimately influence synapse formation and maintenance, which is critical for memory consolidation. Therefore, the RNA-MS data suggests that m^6^A modified *Malat1* contribute to the consolidation of fear-extinction memory through RBP interactions and protein localisation at the synapse.

Finally, upon targeted degradation of m^6^A-*Malat1* at the synapse in the context of fear-extinction learning an impairment in fear-extinction memory is observed. Many of the functional m^6^A studies in the brain have relied on the manipulation of m^6^A methyltransferases and demethylases [13] [9] [14] [15] [70]. Very few studies have looked at the effect of manipulating a single RNA at a specific time and place [71] particularly in the context of fear-extinction learning. Here we used the CIRTS-YTHDF2 single plasmid system with slight [72] modifications to degrade m^6^A-modified *Malat1* at the synapse. The CIRTS-YTHDF2 construct was modified to include a Calm3 intronic region (3’UTR) which can localise the construct to the synapse. As the guide (either control or *Malat1*) is expressed everywhere, only the construct is localised to the synapse, so once the CIRTS-YTHDF2 is at the synapse it can recognise *Malat1* to have its degradative effects. Indeed, binding of YTHDF2 to m^6^A modified RNA has been shown to result in the localisation of RNA from the translatable pool to decay sites (i.e. processing bodies). The carboxy-terminal of YTHDF2 determines selective binding to m^6^A-modified RNA and the amino-terminal domain allows localisation of the entire YTHDF2-RNA complex [73]. As such, the degradation of *m^6^A-Malat1* at the synapse resulted in the impairment of fear-extinction memory. The impairment in fear-extinction memory is likely through the lack of m^6^A-*Malat1* interactions occurring between CYFIP2 and DPYSL2 proteins, thus inhibiting localisation of the proteins to form WAVE complexes and ultimately reducing synapse formation.

In summary, we have the discovered that m^6^A-modified *Malat1* at the synapse is critical for fear-extinction memory, a finding that deepens our understanding of the mechanistic role of m^6^A-modified RNA and its interaction with specific RBPs in subcellular time and space. Together these data support the idea that the state dependent activity of m^6^A-modified RNA is critically important for the consolidation of fear extinction memory.

## Acknowledgments

We thank Dr. Alun Jones from the Mass Spectrometry Facility in the Institute for Molecular Bioscience at the University of Queensland for help with the proteomics experiments and Ms. Rowan Tweedale in the QBI for help in manuscript editing. The authors acknowledge grant support from NIH R01MH109588 (TWB and RCS), NHMRC Ideas Grant (GNT2003414, TWB), NSFC 82001421 [74], the National Institute of General Medical Sciences of the National Institutes of Health NIH (R35 GM119840, B.C.D.). E.L.Z, L.J.L. and S.U.M. are supported by a Westpac Future Leaders Scholarships and the University of Queensland.

## Author Contributions

S.U.M. and T.W.B. conceived this project, together with R.C.S., and led the development and optimization of the protocol, performed experiments, analysed and interpreted data, generated figures, and wrote the paper. H.G., L.J.L., S.U.M., A.P., X.L. W.W. and E.L.Z. assisted with tissue collection, surgery and behavioural experiments. P.R.M. performed the behavioural experiments and analysis. Q.Z. performed bioinformatics analysis. W.S.L. designed the construct, performed the mass spec experiment and helped write the manuscript. All authors discussed results and revised the manuscript.

## Materials and Methods

### Animals

Adult, male C58BL/6 mice (at 9-10 weeks old, 20-25g in weight) were used for experiments. Mice were housed 4 animals per cage on a 12-hour light:dark cycle (lights on 0700h) in a humidity and temperature-controlled vivarium, with rodent chow and water given *ad libitum*. All animal behavioural testing was conducted during the light phase in red-light illuminated testing rooms. Mice were culled by cervical dislocation, the prefrontal cortex (PFC) was dissected from whole brains snap frozen on liquid nitrogen and transferred to −80 storage. All procedures were conducted according to protocols and guidelines approved by the University of Queensland Animal Ethics Committee.

### Synaptosome isolations

Synaptosomes were isolated from PFCs of 60CS 12 RC and 12 extinction trained adult mice (10 weeks of age), using a discontinuous Percoll-sucrose density gradient with modifications. 4 PFCs were dissected and homogenised in ice-cold GM buffer (2.5M sucrose, 1M Tris-HCI) for one synaptosome isolation, leaving 3 RC and 3 EXT samples. A 250uL aliquot of this total cell homogenate was taken and the rest of the homogenate centrifuged at 1,000g for 10 minutes at 4C. After centrifuging, the supernatant was applied to a discontinuous Percoll gradient, comprising of 3%, 10%, 15% and 23% Percoll layers (GE Healthcare) in GM buffer and RNaseOUT (1:1000). The gradient was centrifuged at 18400 rpm for 5 minutes at 4C in a fixed-angle rotor. Four major fractions were separated from the Percoll gradient, where synaptosomes were collected from layers 3 and 4. These layers were pooled for each sample and diluted fourfold with cold GM-buffer and pelleted by centrifuging 14,800 rpm for 30 minutes at 4C. Pellet was transferred to a new tube and stored at −80C until ready for use.

### RNA-immunoprecipitation and mass-spectrometry

Two previously published biotin-labelled RNA oligonucleotides were used in this assay (Dominissini 2012). Streptavidin-conjugated agarose beads were pre-blocked using 1% BSA for 30 minutes at 4C. After 30 minutes, BSA was discarded and beads were resuspended in 1X RNA capture buffer (20mM Tris, pH7.5, 1M NaCl, 1mM EDTA, 1ul RNAse OUT) and 2ug of oligos were added per reaction. Beads and oligos were incubated for 2 hours at 4C. After incubation, beads were washed 3 times with 20mM Tris (pH 7.5), 5-10 minutes per wash. After washing, 1X protein-RNA binding buffer (20 mM Tris, pH7.5, 50 mM NaCl, 2 mM MgCl2, 0.1% Tween-20, 100U/mL RNAseOUT, 1M DTT, 1X PIC) was added to the beads and incubated at 4C until lysates are ready to add to the beads. Nuclear and synaptosome lysates from trained mouse PFC tissue were washed with 1X PBS (8000rpm, 2 minutes, 4C). Lysates were resuspended in RNA immunoprecipitation buffer (150mM KCI, 25mM Tris, pH 7.4, 5mM EDTA, 0.5mM DTT, 0.5% NP40, 100U/mL RNAseOUT, 1M DTT, 1X PIC) and incubated on ice for 30 minutes. After 30 minutes, lysates were centrifuged for 15 minutes at 10 000g (4C). Keep the supernatant and add protein-RNA binding buffer and 30uL of 50% glycerol to each lysate. Now the lysates are ready to add to pre-bound beads and oligos. Nuclear or synaptic tissue lysates were incubated with beads for 2 hours at 4C then UV crosslinked for 2 minutes at 120, 000kJ. After crosslinking, beads were washed 2-3 times with wash buffer (20mM Tris, pH7.5, 10mM NaCl, 0.1% Tween-20, RNAseOUT, 1M DTT, 1X PIC) for 5 minutes per wash. After the last wash, the beads were sent for LC-MS/MS at the Institute of Molecular Biosciences Mass Spectrometry Unit. Three replicates were conducted for each group (RC-control, RC-m^6^A, EXT-control and EXT-m^6^A). This same method was used for the *Malat1* protein pull down assay but this time using probes against *Malat1*. Each group was manually sorted for high confidence proteins (i.e. >2 hits of the same protein in each replicate and the same protein had to be detected in two or more replicates to be included in further analysis). All proteins were analysed by STRING to determine functional protein association networks and clusters.

### Low-input m^6^A-sequencing

m^6^A-immunoprecipitation was performed as previously published by Dominissini et al., (2012), with slight modifications. Up to 100ng of non-synaptic and synaptic RNA was used. RNA was chemically fragmented into approximately 100 nucleotide fragments by 5-minute incubation at 94C in fragmentation buffer (10mM ZnCl2, 10mM Tris-HCI pH 7). The fragmentation was stopped with 0.05M EDTA. Input samples were stored at −80C. m^6^A-enrichment samples were incubated overnight at 4C with 5ug of affinity purified anti-m^6^A polyclonal antibody (Abcam) in IPP buffer (150mM NaCl, 0.1% NP-40, 10mM Tris-HCI pH 7.4). The mixture was then immunoprecipitated by incubation with protein-G beads at 4C for 2 hours. After extensive washing, bound RNA was eluted from beads in 10uL of water. For m^6^A-enrichment of RNA for qRT-PCR, there were four groups of RNA-input, m6A enriched, unbound RNA or IgG control. RNA was not fragmented. An overnight incubation was also conducted with either the anti-m^6^A antibody or IgG control and the supernatant or “unbound” RNA fraction (i.e. the RNA which did not bind to the m^6^A antibody conjugated beads) was also saved. All samples were washed and eluted in 12uL of water and used for cDNA synthesis.

8uL of RNA was used for library generation with the SMARTer Stranded Pico Input Mammalian Takara kit, according to the manufacturer’s instructions. Each library was subjected to AMPure XP (Beckman Coulter) purification to remove primer-dimers and fragments larger than 500bp. The final libraries (size distribution from 200-500bp, with the peak at approximately 400bp), were run on a single-flow cell on Hiseq 4000 (Illumina) for paired-end sequencing.

### m^6^A-sequencing data analysis

Cutadapt was used to trim off low-quality nucleotides (Phred quality <20) and adaptor sequences at the 3’ end. All reads were then aligned to further filter reads mapped to rRNA or Phi X genomes. HiSAT (v2.1.0) was used to align the filtered reads to the mouse genome (mm10). Once aligned, significantly modified transcripts in RC vs. EXT groups were determined using ExomePeak (v2.16.0). All bioinformatic analysis was performed by Dr. Qiongyi Zhao. These transcripts were used in Gene Ontology functional analysis using the DAVID Bioinformatic Resources. Also, the Integrative Genomics Viewer software was used to view m^6^A peaks in each sample.

### Neuronal culture

Cortical neurons from embryonic day 16 (E16) mice were prepared and maintained in neurobasal medium containing 2% B28, 2mM Glutamax, 50U/mL penicillin and 50ug/mL streptomycin. Neurons were transduced with lentivirus at 3 days *in vitro* and harvested for experiments 10 days *in vitro*, after a 4hr, 20mM KCI stimulation.

### fRIP

Formaldehyde RNA immunoprecipitation was performed with modified protocols from [75] and Rinn et al., (2018). Samples were homogenised in PBS and cross-linked using 0.1% formaldehyde (methanol free) for 5 minutes at room temperature. Cross-linking was quenched using 125mM glycine. Samples were centrifuged at 8000rpm for 2 minutes and then washed extensively using PBS and 1X protease inhibitor. Samples were resuspended in 1mL lysis buffer (25mM Tris-HCI pH7.5, 150mM KCI, 5mM EDTA and 0.5% NP-40 with 1:1000 RNaseOUT, 1M DTT and 1X PIC fresh). 56uL of each sample was reserved as the input material (i.e. does not undergo immunoprecipitation) and the rest was split in half (one half undergoes IP with DPYSL2 antibodies and the other half with CYFIP2 antibodies). 5ug of antibody was added to each IP and rotated at 4C for 2 hours. After the 2-hour incubation, 50uL of Protein G beads were added to the antibody-lysate mix and rotated for 1 hour at 4C to isolate protein bound RNA from the beads. Beads were washed with native lysis buffer and resuspended in 56uL of ultrapure water, which was reverse-crosslinked. 33uL of reverse-crosslinking buffer (3X PBS, 6% N-Lauroyl Sarcosine, 30mM EDTA, 15mM DTT, 1:1000 RNaseOUT and 1X PIC) was added to all samples and incubated at 42C for 1 hour then 55C for another hour to separate cross-linked RNA and protein. RNA was extracted using RNAClean XP beads and DNAse treated (using the Zymo DNase enzyme and buffer as per manufacturer’s protocols). RNA was eluted in 11uL of RNAse-free water and used for qRT-PCR.

### RNA extraction

For all RNA samples (from neuronal cultures and PFC tissue) RNA was first extracted using Nucleozol reagent (Invitrogen), as per manufacturer’s protocol. At the phase separation step, where the nuclei and protein are pelleted, the supernatant was combined with an equal volume of 95-100% ethanol and transferred to a RCC5 column part of the RNA Clean and Concentrator kit from Zymo Research and manufacturer’s instructions were followed.

### qRT-PCR

Up to 1ug RNA was used for cDNA synthesis using the Quantitect Reverse transcription Kit (Qiagen). qPCR was performed on a RotorGeneQ (Qiagen) using SYBR-Green Master Mix (Qiagen), with primers for target genes (*Malat1*) on a RotorGeneQ (Qiagen) real-time PCR cycler. All transcript levels were normalised to input RNA using the %input method and each PCR reaction was run in duplicate for each sample.

### Cloning

The CRISPR-Cas-Inspired RNA targeting system (CIRTs) was used to demethylate or degrade modified RNA, because it is a single construct of which the protein component is much smaller than earlier iterations of CRISPR-Cas (Rauch 2019). The pFsy(1.1)GW lentiviral expression vector (addgene #27232), containing the synapsin 1 promoter, was used to make the Syn1-Cirts-Calm3 construct. Upon synaptic activity, it has been demonstrated that STAUFEN binds to the 3’UTR of *Calm3* to mediate localisation of *Calm3* to dendrites [76]. Therefore, after transcription of the components and neuronal activation, the addition of the *Calm3* 3’UTR allows the entire construct to be localised to the synapse. As the guide is expressed everywhere, upon localisation of the construct to the synapse, the construct can be used to degrade/demethylate m^6^A-*Malat1* at the synapse.

The CIRTs pin nuclease construct, containing a C-terminal Flag tag and nuclear export signal (NES), was a kind gift from Bryan Dickinson group (Rauch et. al. 2019). The CIRTs cassette was PCR amplified and cloned into AgeI and XbaI site. A NheI site was generated upstream of XbaI using PCR primer. The Calm3 dendritic localization sequence was PCR amplified from psiCheck2-Calm3 (a kind gift from Michael Kiebler’s lab), and cloned into the NheI and XbaI site. Finally, the U6-gRNA scaffold was PCR amplified from CIRTS pin nuclease construct and inserted in the XbaI site to make the final construct.

For the CIRTs YTHDF2 Calm3 plasmid, the above Syn1-CIRTs-Calm3 construct was digested using AgeI and NheI. The original CIRTs-6: B-defensin 3-TBP6.7-YTHDF2 plasmid (# 132544) was purchased from Addgene and the YTHDF2 domain was PCR amplified and cloned into the AgeI and NheI sites. The CIRTs FTO plasmid was constructed similarly, using the full-length FTO sequence synthesised by IDT. GFP was PCR amplified and cloned into AgeI site along with a 2A peptide signal.

Finally, to clone the gRNAs into the plasmids, DNA fragments were synthesised by IDT. Three gRNAs against *Malat1* (corresponding to m^6^A peaks identified in Fig. 2) or a control guide were cloned into the constructs to generate three individual *Malat1* targeting constructs and one control construct. Each guide had the U6 promoter (with the XbaI site), followed by TBP fold (RNA hairpin binding domain), guide sequence and overhang with a EcoRI site. Guides were PCR amplified and cloned into the plasmid using XbaI and EcoRI sites.

### Lentiviral Surgery

Lentivirus was prepared as previously described (Li et al., 2014). Double cannulae (PlasticsOne) were implanted in the anterior posterior plane, along the midline into the infralimbic prefrontal cortex (ILPFC), a minimum of 7 days prior to viral infusions. Injection coordinates were centred at +1.85mm in the anterior posterior (AP) plane and −2.5mm in the dorsal-ventral (DV) plane. A total of 2uL of lentivirus was introduced via two injections, delivered at a rate of 0.1uL/min, 48 hours apart. 24-hours prior to lentiviral infusions animals were fear conditioned. One-week after lentiviral infusions mice were extinction trained.

### Behavioural tests

Two contexts (A and B) were used for all behavioural fear testing. Both conditioning chambers (Coulbourn Instruments) had two transparent walls and two stainless steel walls with a steel grid floors (3.2mm in diameter, 8mm centres); however, the grid floors in context B were covered by flat white plastic transparent surface. This surface is used to minimise context generalisation. Digital cameras were mounted in the ceiling of each chamber and connected via a quad processor for automated scoring in a freezing measurement program (FreezeFrame). Fear conditioning (context A) was performed with a spray of lemon-alcohol (5% lemon and 10% alcohol). The fear-conditioning protocol started with a 120 sec pre-fear conditioning incubation which was followed by three pairings of a 120 sec, 80dB white noise conditioned stimulus (CS) with a 1 sec, 0.7mA foot shock (US). Based on fear conditioned freezing percentages mice were matched into equivalent treatment groups (Malat1 virus or control virus). For extinction (context B) which was performed with a spray of vinegar, mice were again allowed to acclimate for 2 minutes and then extinction training involved 60 nonreinforced 120sec CS presentations (a 60CS training protocol). For the behaviour control experiments, animals did not receive the CS (this group is known as the retention control - RC). Memory was tested by returning animals to context B (24 hours apart) with a 3CS-only presentation protocol. In these tests, the CS was presented but no foot shock (US) was delivered. Memory was calculated as the percentage of time spent freezing during the tests.

For tissue collection for synaptosome isolations, 24 mice were fear-conditioned then 12 were taken as RC and 12 were 60CS extinction trained, 24 hours later. Immediately after extinction training, mice were sacrificed and PFC tissue was snap frozen for synaptosome isolations. Each replicate had non-synaptic and/or synaptic fraction for subsequent experiments.

### Statistical analysis

All statistical analyses were performed using Prism 9. Two-tailed unpaired Student’s t-test was used comparison between -KCI and +KCI groups in m^6^A enriched, unbound and IgG control fractions as well as CIRTs Control and *Malat1* groups. One-way or two-way ANOVA was chosen for multiple comsparisons where appropriate. All post-hoc analysis was performed using Dunnet’s multiple comparison test. Error bars represent SEM where significance was taken as p<0.05.

